# Trajectories of Alcohol Use Initiation and Risk to Develop an Alcohol Use Disorder During Adolescence: A Role for Stress and Amygdala Activity

**DOI:** 10.1101/178251

**Authors:** Nourhan M. Elsayed, M. Justin Kim, Kristina M. Fields, Rene L. Olvera, Ahmad R. Hariri, Douglas E. Williamson

**Affiliations:** Translational Center for Stress-Related Disorders, Department of Psychiatry and Behavioral Sciences, Duke University School of Medicine, Durham, NC; Laboratory of NeuroGenetics, Department of Psychology and Neuroscience, Duke University, Durham, NC; Department of Psychiatry, University of Texas Health Center, San Antonio, TX; Research Division, Durham VA Medical Center, Durham, NC

## Abstract

Early alcohol use initiation predicts onset of alcohol use disorders (AUD) in adulthood. However, little is known about developmental trajectories of alcohol use initiation or their putative biological and environmental correlates. Here we report the results of latent class analyses identifying two trajectories of alcohol use initiation in a prospective study of adolescents selected for the presence or absence of familial risk for depression at baseline. The latent class analyses identified two distinct patterns of initiation: early initiators (EI; n=32) who reported greater baseline alcohol use (M = 1.12, SE = .35) and exhibited a more rapid rate of change in use between baseline and each follow up wave (M = 4.43, SE = .94); and in contrast, late initiators (LI; n=298) who reported lower baseline use (M = .23, SE = .03) and exhibited a slower rate of change between baseline and each of the subsequent follow up waves (M = .12, SE = .03). Early initiators had more positive expectancies regarding alcohol (p = .002 – p = .005) and reported higher levels of stressful life events during the year prior to baseline assessment (p = .001). Additionally, fMRI analyses revealed that EI displayed heightened threat-related amygdala activity at baseline compared to LI (p = .001), but no differences in reward-related ventral striatum activity. Lastly survival analyses revealed that EI initiators were 6.7 times more likely to develop an AUD by age 19 when compared to the LI (p = .005). These patterns, which were independent of broad familial risk for depression, suggest that early initiation of alcohol use during adolescence associated with later risk for AUD is reflected in both higher levels of stressful life events and higher neural reactivity to threat, the combination of which may inform ongoing efforts to prevent persistent dysfunction.

## Introduction

Alcohol use disorder (AUD) is a significant public health problem in the United States with approximately one-third of the population experiencing a life-time episode of AUD^1^. Recent data from the Monitoring the Future Study showed that as many as 23% of 8^th^ graders have used alcohol at some point in their lifetime, and nearly 9% reported being drunk at least once with these estimates increasing to 61% and 23% by 12^th^ grade, respectively ^2^. While experimentation with alcohol during adolescence is common, earlier age of alcohol use initiation is one of the strongest predictors of lifetime AUD diagnosis and predicts a more severe chronic course, even after controlling for family and demographic factors ^3–8^. Given the public health significance of AUD, and the critical developmental period of adolescence during which alcohol use is typically initiated, it is important to identify the risk factors that are associated with early alcohol use initiation and those associated with progression to AUD ^7,9,10^.

Several demographic and behavioral risk factors have been associated with earlier initiation of alcohol use including being male and white ^11^, and having more positive expectancies regarding the effects of alcohol ^3,12,13^. Other risk factors for early alcohol use initiation include having parents who consume more alcohol and are accepting of teen drinking ^3,10,11^, deviant peer relations, associating with peers who consume alcohol, and having positive perceptions about the number of friends who drink ^3,7,11,14,15^. Importantly, externalizing symptomology is often considered a risk factor and key precursor to earlier alcohol use initiation, ^9,13–16^ and is central in Cloninger’s Type II classification of alcoholism characterized by impulsivity and earlier age of AUD ^17^.

In addition to the classic externalizing disorders and deviant peer group pathways to alcohol use, a relationship between early onset AUD and negative affect has also been observed. In a systematic review, Hussong et al., (2017) found that clinical depression and depressive symptoms predict earlier onset of alcohol use ^18–20^ and are indicators of problem drinking ^21^ as well as overall use ^16^. One explanation for these findings is that alcohol use represents a form of self-medication through which those with high negative affect can reduce their experience of symptoms ^22^. Alternatively, some suggest that alcohol use is associated with negative reinforcement (e.g., drinking to alleviate negative affect) that subsequently and paradoxically leads to the experience of negative affect ^15,23^. Moreover, offspring of depressed parents have been shown to be at increased risk not only to develop depression but also a substance use disorder (primarily alcohol) ^24,25^.

Recent research has reported the longitudinal course of alcohol use during adolescence and suggests several courses can be identified; these have been commonly identified as a nonuser/stable low-use course, a chronic or high-use course, a course characterized by maturing out of drinking, and a later onset or increasing course ^23,26–28^. Critically, these trajectories are sensitive to which indices, or combination of indices were used to ascertain the trajectories, ranging from measuring non-specific alcohol involvement (e.g., alcohol consequences, alcohol dependence) to measuring alcohol quantity– frequency ^27^. Extending from this approach, within a longitudinal framework, it is important to examine trajectories of alcohol use initiation and their correlates as they impact risk for AUD. It is likewise important to model the complexities of alcohol use initiation so that specific risk factors can be evaluated ^23,28,29^.

Here, we first sought to identify patterns of initiation and use in a longitudinal cohort of adolescents at high and low familial risk for depression using trajectory analyses. Next, informed by the results of the trajectory analyses, we sought to identify how trajectories of use differed with respect to putative behavioral risk factors as well as both distal and proximal stressors. Lastly, we examined how the trajectories differed with respect to two neural markers of risk for problem drinking: relatively increased threat-related amygdala activity and decreased reward-related ventral striatum activity in the baseline assessment of adolescents^30,31^. Based on prior research, we hypothesized that increased exposure to stress, enriched familial history for depression, and increased threat-related amygdala reactivity would be associated with greater alcohol use and possibly an early initiation of alcohol use.

## Materials and Methods

### Sample

Participants in this study were recruited as part of the Teen Alcohol Outcomes Study (TAOS); sampling procedures for TAOS have been described previously ^32–36^. Briefly, 1,089 youth between the ages of 12 years 0 months and 14 years 11 months were randomly contacted within a 30-mile radius of the University of Texas Health Center at San Antonio and screened for being at high and low familial risk for depression. Adolescents were determined to be at high familial risk for depression if they had at least one first-degree and one second-degree relative with a lifetime history of major depression. Adolescents were identified as being at low-risk for depression if they had no first-degree and minimal second-degree relatives (< 20%) with a lifetime history of depression ^25^.

A total of 330 participants were recruited into the study; 164 to the high-risk and 166 to the low-risk group. In addition to meeting criteria for familial risk, adolescents were excluded if they met criteria for any psychiatric diagnoses at the baseline assessment (with the exception of anxiety in the high-risk group) including externalizing (e.g. Conduct Disorder, ADHD), or had already binge drank according to NIAAA criteria^37^. Participants were re-assessed annually with diagnostic interviews, and self-report measures of behavior including mood, anxiety, stress, and substance use. All 330 adolescents were followed at least once post-baseline with an overall mean number of 3.86 ± 1.41 assessments. The study was approved by the Institutional Review Board at both the University of Texas Health Sciences Center at San Antonio and Duke University.

### Childhood Trauma Questionnaire

The Childhood Trauma Questionnaire (CTQ) is a 28-item self-report measure and was used to assess exposure to five different types of childhood trauma: emotional, physical, sexual abuse, emotional and physical neglect^38^. CTQ total score reflects a variety of forms of maltreatment and has good reliability ^39^.

### Stressful Life Events Schedule

The Stressful Life Events Schedule (SLES) was used to assessed the occurrence of stressful life events during the past year ^40^. The use of the SLES and the scoring in this study have been described previously ^32–34^. Briefly, the SLES assesses the presence and occurrence of age appropriate stressors in children and adolescents across several domains (e.g., family, friends, school). Each stressor is given a subjective stress and objective stress rating by a consensus panel.

### Mood and Feelings Questionnaire

Depressive symptoms from the past two weeks were assessed with the 32-item child-report and parent-report version of the Mood and Feelings Questionnaire (MFQ) ^41^. Higher total scores on the MFQ reflect a greater likelihood for the diagnosis of a depressive disorder; the MFQ has been shown to have good reliability and validity^42,43^.

### Youth Self Report, Child Behavior Checklist

Depressive symptoms were further assessed using the Affective Problems, Anxious/Depressed, Withdrawn/Depressed and Anxious Problems subscales of the 120-item Child Behavior Checklist (CBCL) and the Youth Self Report (YSR) ^44,45^. The CBCL and YSR have been found to be valid screens for affective and anxiety disorders^46^.

### Kiddie Schedule for Affective Disorders and Schizophrenia Present and Lifetime version (K-SADS-PL)

Lifetime psychiatric disorders according to *DSM-IV-TR* was assessed using the Kiddie Schedule for Affective Disorders and Schizophrenia for School-Age Children, Present and Lifetime Present Episode version (K-SADS-PL) ^47^. The child and parent/guardian served as informants and summary symptom assessments based on each informant were made by the clinical interviewer. At follow-up, the KSADS-PL was used to assess the onset of an AUD including alcohol abuse or dependence. The K-SADS-PL has been used as a gold standard for the assessment of psychiatric disorders in children and adolescents^42,48^.

### Substance Use Questionnaire

The Substance Use Questionnaire (SUQ) was administered at baseline and all follow-up assessments ^49^. The SUQ’s first item asks: “In the past 12 months, how often did you drink beer, wine, wine coolers or liquor?”. The 11-point scale contained options from 0 - (Not at all) to 11 - (Several times a day). The second item asked participants: “Think of all the times you have had a drink in the past 12 months. How much did you usually drink each time?”. Responses ranged from 0 - (I didn’t drink in the past 12 months) to 13 - (More than 25 drinks). Dimensional scaling of frequency of use (over the past year) and amount of use were multiplied to calculate a “use metric” consistent with our prior work ^33^. A score of 1 on this metric is indicative of consuming approximately less than one can or glass of alcohol 1-3 times a year; a score of 10 on this metric is indicative of consuming approximately one drink 2-3 times a month or having four drinks 4-7 times a year; a score of 20 on this metric is indicative of consuming approximately four drinks once a month or having three drinks 2-3 times a month ^33^.

### The Alcohol Expectancy Questionnaire

The Alcohol Expectancy Questionnaire (AEQ) was used to measure alcohol expectancies^50^. The version of the AEQ used in the present study included 68 statements regarding the effects of alcohol with a true– false response. The Cognitive and Motor Impairment, Global Positive Changes, Increased Arousal, Relaxation and Tension Reduction, Sexual Enhancement, Changes in Social Behavior and Improved Cognitive and Motor Abilities subscales were scored, with higher scores indicating more positive alcohol expectancies.

### fMRI Tasks

Participants performed an emotional face matching task associated with robust threat-related amygdala activity. Here we examined amygdala activity to general threat-related signals as represented by angry and fearful facial expressions, respectively, as well as amygdala activity to each expression independently. The latter allows for modeling of amygdala activity to both explicit, interpersonal threat in the form of angry facial expressions and implicit, environmental threat in the form of fearful facial expressions^51^. Participants also performed a card guessing task with monetary incentive to elicit reward-related ventral striatum activity. These tasks were administered to all participants at baseline. Further details are presented in the supplementary materials. Both tasks have been used extensively including prior studies reporting data from the present sample ^32,34–36,52^.

### Growth Mixture Modeling

Growth Mixture Models (GMM) were used to identify groups of adolescents with similar intercepts and courses of alcohol use. Per Ram & Grimm’s suggestion a baseline growth model was used to establish that a linear trajectory is the best representation of change in the longitudinal data ^53^. Subsequently, we proceeded to the selection of an optimal number of class trajectories by following an iterative sequence ^53–55^.

As part of this sequence, model fit was evaluated through comparisons of the Lo-Mendall-Rubin-Adjusted Likelihood Ratio Test p-value, Bayesian Information Criterion (BIC) ^53,54,56,57^, posterior probabilities, entropy, and total group size ^47,51^. Model fit of a “k class” model (e.g., a 4-class model) was compared to model fit of a “k-1 class” model (e.g., a 3-class model). This process concluded and the optimal number of classes was determined when the likelihood-ratios tests were no longer significant, when over-extraction is evident, or when a class model fails to converge ^59^. Additionally, for each number of classes, we specified models that allowed for expected parameter differences within class. All models were run with 400 random starts.

Once the group trajectories were identified, each case was assigned to a class based on their most likely class membership. Participants were grouped by Modal assignment given the value of entropy > .9 for the final identified class structure ^60^. Between trajectory group differences were examined using Welch’s F ratios to adjust for differences in the homogeneity of variance. Bonferroni adjusted alpha levels were used to conservatively adjust for the number of differences examined.

All LGCM analyses were fit using Mplus version 8.0 ^56^. Given the longitudinal nature of the data, some cases were missing due to attrition and thus a full-information maximum-likelihood estimator (FIML) was used, as these models assume the data are missing at random and produce estimates based on all available data ^56^. Descriptive and behavioral analyses were completed using SPSS (IBM Corporation, Armonk, NY, United States) and R version 3.3.3 (2017-03-06). More information regarding the LCGM can be found in the supplementary materials section of this manuscript.

### fMRI Statistical Analyses

Blood-oxygen-level-dependent (BOLD) fMRI data were analyzed using SPM8. Further details on quality control procedures and extraction of BOLD parameter estimates from task-related functional clusters are provided in the supplementary materials. A 2 (alcohol initiation) × 2 (wave) analysis of variance (ANOVA) model was used to examine the fMRI data for each experiment (amygdala and VS, respectively) predicted by class. Welch’s t-test was used for post hoc comparisons following the outcomes of the omnibus ANOVA tests.

### Survival Analyses

Survival analysis examined the relationship between group membership and age at onset of AUD. “Survival” is defined as the absence of alcohol use disorder at the time of last assessment; “death” is the onset of a disorder. The analyses examined the age at onset for adolescents developing an AUD and age at last assessment was used to censor adolescents not developing an AUD. Kaplan–Meier estimator was used to estimate the unadjusted survival function across survey waves. Time-dependent Cox proportional hazards models were used to estimate the unadjusted and adjusted hazard ratios (HRs) for onset of alcohol use disorder at the current wave in relation to LCGM-identified group membership (EI and LI). Survival analyses were performed using SPSS statistic software (SPSS Inc., USA).

## Results

### Identification of Alcohol Use Initiation Trajectories

The models described in the methods were fit to the longitudinal alcohol use data presented in Supplementary Figure 1. For the two and three-class models, we encountered convergence issues including variance estimates that were negative. In these cases, we fixed covariance’s to zero and re-ran the models ^53^ (See Supplementary Table 1). The models began to fail to converge with the three class models suggesting that a model with more than three classes was unlikely, thus we ceased to run models with more than three classes (See Supplementary Table 1).

**Figure 1.**
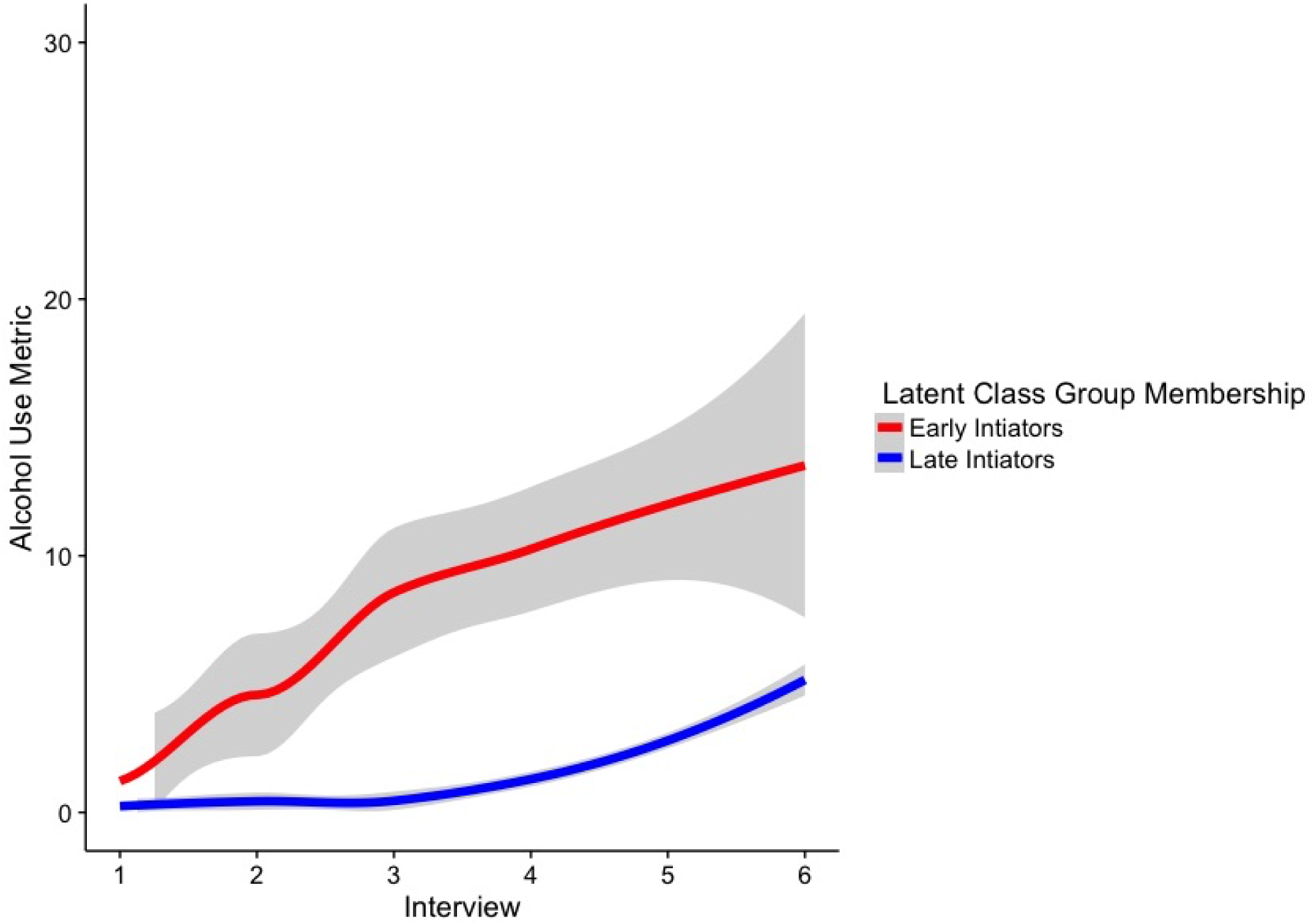
Class-specific alcohol use trajectories based on estimated means (± 95% CI).

The model which estimated differences in means between two classes and which restricted the intercept and slope variance for class two to zero, but which allowed these variances to be freely estimated in class one was the best fitting model. This model had the lowest information criteria (i.e., AIC, BIC, ABIC) value, had a value of 0.92 for entropy and all three Likelihood Ratio Tests supported the use of this 2-class model. After selecting this model, we evaluated the differences in parameters between the two groups (see Supplementary Table 2).

Group one contained 32 adolescents whose trajectories were labeled the early initiators (EI) type with a progressive increase in alcohol assumption over time (see Figure 1). The second group, containing 298 participants were classified as later initiators (LI) type, with relatively no alcohol use at baseline and a slower increase in either initiation of alcohol use or continued absence across adolescence.

Across the first five waves of the study, the EI had greater alcohol use levels. At the sixth study assessment the two groups, had comparable levels of alcohol consumption (see Figure 1 and Supplementary Table 3).

### Demographic and Behavioral Correlates of Alcohol Initiation

The two groups were similar in age, ethnicity, gender, number of interviews, and familial risk for depression (see Table 1). In addition to comparing the two groups on demographic characteristics, we examined baseline behavioral characteristics between the two groups. After correction for multiple comparisons, the AEQ - Increased Arousal sub-scale (p = .002), the AEQ - Social Behavior sub-scale (p = .002) and overall SLES objective stress (p = .001) were all significantly higher in the EI group. The differences between the two groups remained through follow-up two, however, they were not consistently repeated throughout the remaining 4 time points (see Table 2).

**Table 1.** Participant demographic descriptive statistics

|  | Early Initiators (EI) | Late Initiators (LI) | p |
| --- | --- | --- | --- |
| Age | 13.32 (0.87) | 13.42 (0.97) | 0.54 |
| Gender | M = 14 | M= 149 | 0.50 |
|  | F = 18 | F = 149 |  |
| Race | White = 19 | White = 167 | 0.32 |
|  | Other =13 | Other = 131 |  |
| Familial Risk for Affective Disorders | High = 16 | High = 146 | 0.91 |
|  | Low = 16 | Low = 152 |  |

**Table 2.** Phenotypic differences between early and late alcohol initiators

|  |  | Early Initiators<br>(EI) | Late Initiators (LI) | p |
| --- | --- | --- | --- | --- |
| AEQ | Cognitive and Motor Impairment | 2.72 (0.68) | 2.69 (0.65) | 0.808 |
| AEQ | Global Positive Changes | 0.75 (0.76) | 0.52 (0.86) | 0.116 |
| AEQ | Increased Arousal | 1.88 (0.94) | 1.29 (1.06) | 0.002* |
| AEQ | Relaxation and Tension Reduction | 2.13 (1.01) | 1.72 (1.20) | 0.043 |
| AEQ | Sexual Enhancement | 1.31 (1.04) | 1.22 (0.98) | 0.656 |
| AEQ | Changes in Social Behavior | 0.75 (0.80) | 0.27 (0.53) | 0.002* |
| AEQ | Improved Cognitive and Motor Abilities | 0.16 (0.37) | 0.11 (0.33) | 0.526 |
| CBCL | Anxiety Problems | 0.6 (1.33) | 0.73 (1.10) | 0.607 |
| CBCL | Anxious/Depressed | 1.65 (2.12) | 1.78 (1.92) | 0.73 |
| CBCL | Withdrawn/Depressed | 0.9 (1.40) | 1.12 (1.82) | 0.438 |
| CBCL | Affective Problems | 1.13 (2.26) | 0.88 (1.62) | 0.548 |
| MFQ_C | Summary Score Child Report | 11.34 (11.17) | 9.41 (8.19) | 0.349 |
| MFQ_P | Summary Score Parent Report | 4.53 (5.82) | 3.1 (5.19) | 0.192 |
| SCARED_C | Summary Score Child Report | 13.28 (7.97) | 16.13 (10.53) | 0.071 |
| SCARED_P | Summary Score Parent Report | 10.72 (19.25) | 6.41 (6.20) | 0.217 |
| YSR | Anxious/Depressed | 3.18 (2.33) | 3.79 (3.26) | 0.208 |
| YSR | Withdrawn/Depressed | 2 (1.98) | 2.69 (2.20) | 0.087 |
| YSR | Affective Problems | 2.83 (2.56) | 2.81 (2.73) | 0.974 |
| YSR | Anxiety Problems | 1.89 (1.42) | 2.13 (1.93) | 0.432 |
| SLES | Objective Stress Rating | 12.42 (6.84) | 7.94 (5.08) | 0.001* |
| CTQ | Total Score | 35.4 (10.52) | 32.89 (6.93) | 0.212 |

\* Significant at $p < .05$ after Bonferroni correction

### fMRI Correlates of Alcohol Initiation

#### Amygdala Activity in Early vs. Late Initiators

Overall, the task produced robust amygdala activity to all faces > geometric shapes (left amygdala, MNI −22, −2, −18, t_(230)_ =15.69, k = 172 voxels, p_FWE-corrected_ < .05; right amygdala, MNI 22, −4, −18, t_(230)_ =18.96, k = 231 voxels, p_FWE-corrected_ < .05; similar results were observed for fearful faces > geometric shapes and angry faces > geometric shapes). Analyses revealed significant differences in between the EI versus LI at baseline (n = 231; 22 EI, 209 LI; Welch’s t_(29.21)_ = 3.72, p = .0008) (see Figure 2a & 2b).

**Figure 2.**
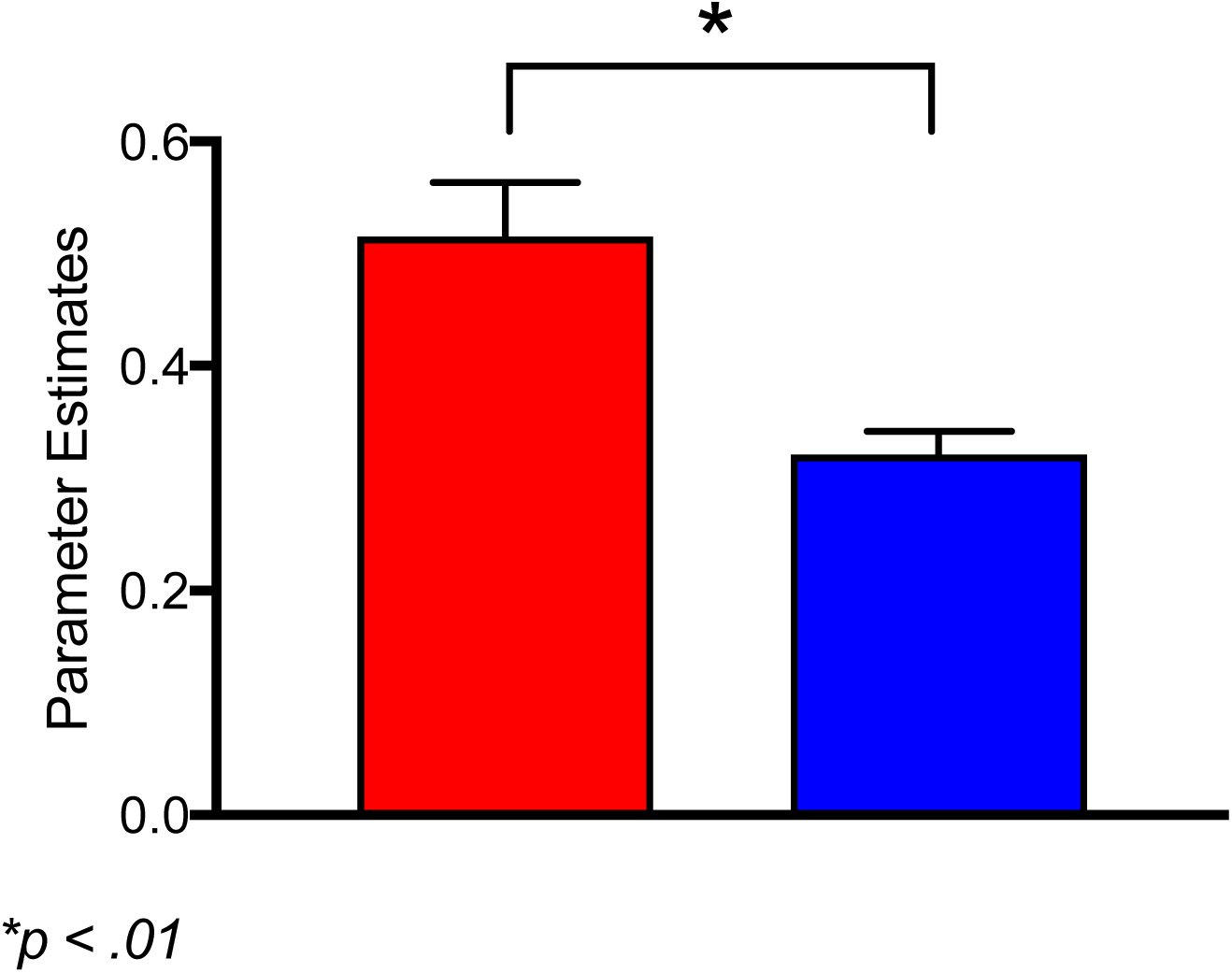
**Figure 2a (**above**) and 2b (**below). Amygdala activity associated with implicit, interpersonal threat (i.e., fearful facial expressions) as a function of early alcohol use initiation at baseline and follow up. Analyses revealed significant differences in between the EI (**red**) versus late LI (**blue**) at baseline (*p = .0008.)* with EI showing more activation in response to fearful faces vs shapes (below).

#### Ventral Striatum Activity in Early vs. Late Initiators

The task elicited significant activity in the left VS to positive > negative feedback (MNI −14, 14, −12, t_(212)_ =4.39, k = 51 voxels, p_FWE-corrected_ < .05. There were no significant differences in VS activity to positive > negative feedback (or any other contrast) between the EI vs. LI (n = 213; 20 EI, 193 LI; all ps > .05) (see supplement Figure 2a & 2b).

### New-Onset of an Alcohol Use Disorder

A total of 9 (2.73%) of the 329 adolescents in the study developed an AUD based on DSM-IV TR criteria. Alcohol use disorder occurred in 5 out of 298 late initiators (1.67%) of and 4 out of 32 early initiators (12.5%). Kaplan–Meier survival estimates examined the cumulative probability of developing an AUD in EI compared with LI adolescents. This survival difference was statistically significant using the log-rank test (χ^2^ = 10. 73, df = 1, p = .001) (see Figure 4). Similarly, Cox proportional hazards regression showed the EI to be 6.7 times more likely to develop an AUD, [95% confidence interval (CI): 1.79–24.94, p = .005] compared with LI (see Figure 3).

**Figure 3.**
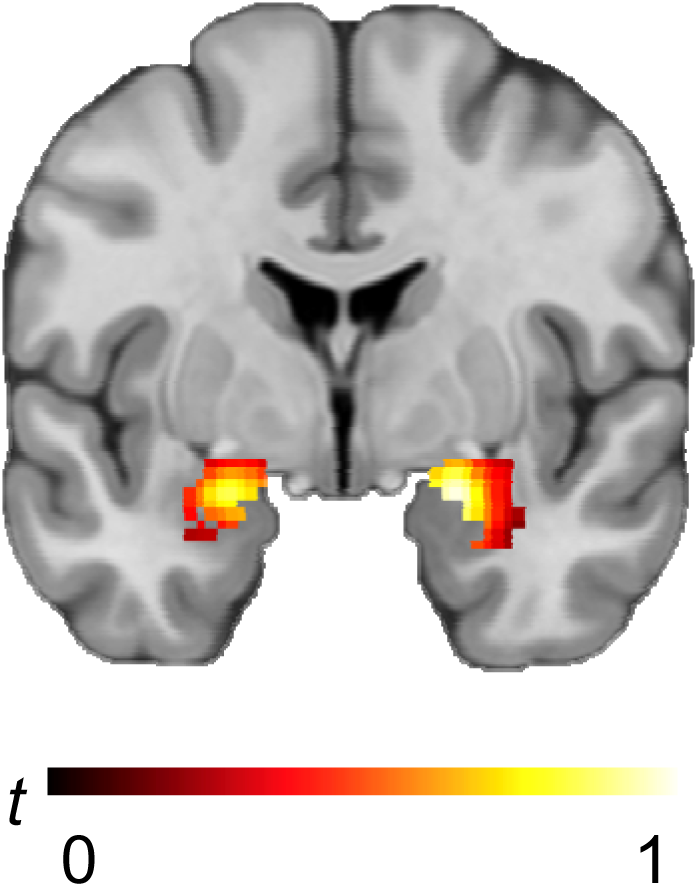
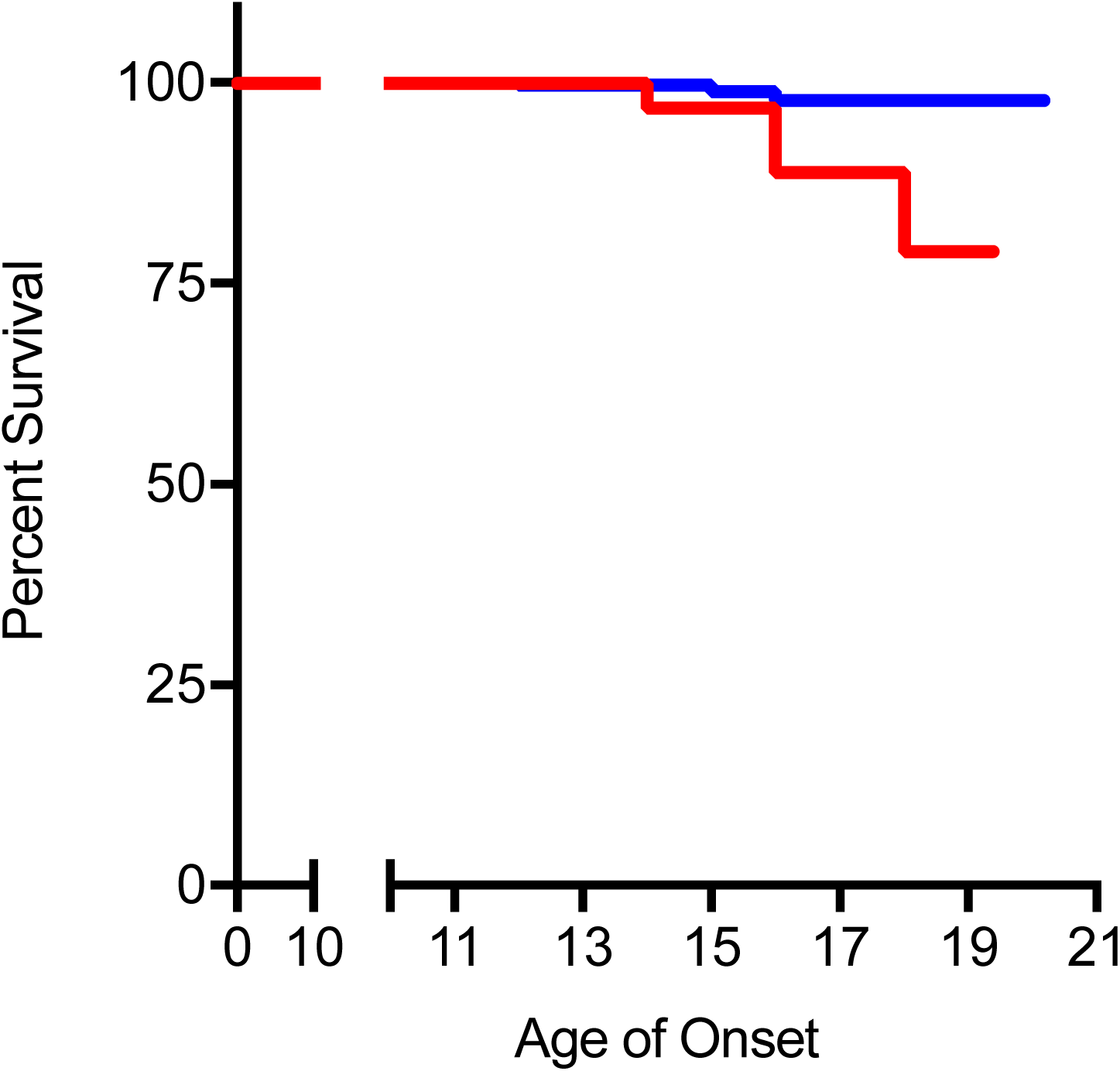
Kaplan–Meier survival curve to incidence of substance use disorder. During the follow-up period, alcohol use disorder diagnosis occurred in 5 of the 298 LI **(blue**) and four of the 32 EI (**red**). On survival analysis, the cumulative probability of developing substance use disorder for early initiator adolescents was higher than those for later initiator adolescents.

## Discussion

In this report, we examined trajectories and correlates of alcohol use initiation in a sample of adolescents enriched for familial risk for depression who were primarily alcohol naïve at their baseline assessment. We identified two classes of alcohol initiators; early (EI) and late (LI). EI had higher amygdala activity to implicit, environmental signals of threat in the form of fearful facial expressions, had more positive expectancies regarding alcohol consumption, and had higher recent stress at baseline. Moreover, EI had greater quantities of alcohol consumption, a more rapid acceleration in use, and were at increased risk for developing an initial AUD. Our findings suggest the importance of higher levels of stress and greater amygdala activity among adolescents who will initiate alcohol use earlier, and add to our prior observations in a subset of these adolescents of increased regional cerebral blood flow in mesolimbic regions associated with current and future alcohol use initiation ^33^. Our findings are consistent with those of Nikolova et al. (2016), who reported that higher amygdala activity to implicit, environmental threat specifically predicted future problem drinking in response to stressful life events as well as AUD in university students^30^. However, the effect of higher amygdala activity observed in their study was expressed as problem drinking and AUD only in participants with relatively low reward-related VS activity. We observed no such moderating effects of VS activity in our analyses, nor did we observe any primary differences in VS activity between EI and LI groups.

Our findings are contrary to several prior reports showing that amygdala hypoactivity is associated with problem drinking and heightened familial risk for AUD ^56–59^. There are several possible contributors to this divergence. First, prior studies did not isolate amygdala activity to different forms of threat and thus, risk-related hypoactivity may not apply to implicit, environmental threat. Second, the extant literature captures a broad and not necessarily overlapping age range of participants. Thus, risk may be associated with relative hyper- or hypoactivity across different developmental windows as has been observed in studies of mood and anxiety ^61,62^. Lastly, the contribution of amygdala hypoactivity to risk may be conditional on the dynamics of other neural circuits. Interestingly, Nikolova et al. not only found evidence for increased stress-related problem drinking in participants with relatively high amygdala and low VS activity, but also in participants with the opposite pattern of relatively low amygdala and high VS activity ^63^. The authors demonstrated that the former risk pattern is associated with higher negative affect and possible problem drinking as a form of coping, while the latter pattern is associated with higher impulsivity and possible problem drinking as a form of disinhibition and poor decision making. This may be consistent with the observation of amygdala hypoactivity in prior studies of participants with familial risk for an AUD where impulsivity is a likely factor, and our observation of hyperactivity in participants with familial risk for depression where negative affect is a likely factor. However, we observed no significant moderating effects of familial risk for depression in our analyses.

We observed that early alcohol use initiation occurred equally in adolescents at high and low familial risk for depression, and that the increased risk for AUD onset in the early initiators was independent of familial risk for depression, internalizing, and externalizing symptomology. By excluding adolescents with externalizing symptomology and enriching the sample for familial risk for depression, we aimed to examine an affective pathway for alcohol initiation and AUD during adolescence. This was motivated by our interest to link prior research that established a relationship between familial risk for depression and lifetime AUD ^62^ with that showing early alcohol use initiation increases risk for lifetime AUD ^9–11,65–69^. However, we did not find independent or additive effects of familial risk for depression or internalizing symptomology on early alcohol use initiation or AUD onset. It is worth noting that the average age at our final follow up was 18.4 years, while the reported average age of onset of AUD of 22 to 24 years ^70^. Therefore, it is possible that the link between depression and AUD will emerge as these participants enter young adulthood, at which time we will be able to determine whether familial risk for depression plays a role in the onset of AUD. This likelihood is underscored by prior research showing that alcohol primarily serves as a negative reinforcement ^15^ and that negative affect precedes the onset of AUD.

Although our data do not support a relationship between familial risk for depression and alcohol use initiation, they do suggest a pathway between early alcohol use initiation and recent life stress. These results converge with prior research linking age at first drink with stress-reactive drinking ^66^. Others have found that after controlling for externalizing symptomology and other sociodemographic risk factors, high stress levels in the past year are associated with greater increase in consumption of alcohol only amongst individuals who initiated alcohol use early ^65,66,68^. Dawson suggests that this relationship is because individuals who initiate alcohol use earlier use adverse coping styles that result in excess consumption thus put them at risk for developing alcohol use disorders ^5,65,66,71,72^. In our cohort, the differences in stress between groups is ameliorated by the second interview wave suggesting that an adolescent may use alcohol as a method of coping and may continue to use alcohol after the stress has subsided. This finding is also in accordance with the literature reporting that age at first drink moderates the impact of stressful life events on later drinking behavior ^71^. The importance of recent stress is underscored by the higher amygdala activity to fearful facial expressions observed in the EI group, which may contribute to increased negative affect in response to these stressors ^35^.

The finding that early initiators escalate their drinking more rapidly and consume more alcohol is consistent in part with prior research showing that escalation during adolescence leads to more negative alcohol dependence outcomes like increased risk of alcohol related problems and alcohol dependence ^23,73^. Further, our observation that EIs were six times more likely to have an AUD by aged 19 years has similarly been observed across numerous prior studies ^5,65,74,75^. Our findings also corroborate prior literature, which has shown that there is more than one pattern of alcohol use initiation in adolescence ^19,23,26,28^. However, our findings diverge in that we identify two unique trajectories of alcohol use initiation, whereas other literature has reported on as many as five trajectories of alcohol use initiation ^19,23,26,28^. Our identification of two trajectories of alcohol initiation may reflect the sampling strategy used that excluded adolescents with internalizing and externalizing symptomology disorders and was enriched for familial risk for depression. In addition, we derived an alcohol use metric based on the quantity and frequency of alcohol use; other research has suggested that there is variability in the number of latent groups identified because there is differences in metrics of use reported^27^.

Our study is not without limitations that can be considered in future research. First, a limited number of questions capturing quantity and frequency of alcohol use were used. It is possible that a fuller spectrum of alcohol use questions could have identified a different pattern of alcohol initiation in this cohort. Second, our overall sample of 329 adolescents while large for a neuroimaging study, is relatively small for GMM analyses which depend on large samples. However, our model fit statistics suggest that while our sample is small, our overall model fit to these data are excellent. Third, only a small number of adolescents were followed long enough to document the initial onset of an alcohol use disorder. Therefore, our analyses examining the contribution of early alcohol initiation and amygdala reactivity to initial AUD onset must be viewed with caution until more of the sample has been followed into young adulthood.

